# Striatal integration of inverse dopamine and serotonin signals gates learning

**DOI:** 10.1101/2023.06.14.544997

**Authors:** Daniel F. Cardozo Pinto, Matthew B. Pomrenze, Michaela Y. Guo, Brandon S. Bentzley, Neir Eshel, Robert C. Malenka

**Affiliations:** Nancy Pritzker Laboratory, Dept. of Psychiatry and Behavioral Sciences, Stanford University School of Medicine, Stanford, CA 94305; Magnus Medical, Burlingame, CA 94010

## Abstract

The neuromodulators dopamine (DA) and serotonin (5-hydroxytryptamine; 5HT) are powerful regulators of associative learning^1–9^. Similarities in the activity and connectivity of these neuromodulatory systems have inspired competing models of how DA and 5HT interact to drive the formation of new associations^10–13^. However, these hypotheses have yet to be tested directly because it has not been possible to precisely interrogate and manipulate multiple neuromodulatory systems in a single subject. Here, we establish a double transgenic mouse model enabling simultaneous genetic access to the brain’s DA and 5HT systems. Anterograde axon tracing revealed the nucleus accumbens (NAc) to be a putative hotspot for the integration of convergent DA and 5HT signals. Simultaneous recordings of DA and 5HT input activity in the NAc posterior medial shell revealed that DA axons are excited by rewards while 5HT axons are inhibited. Optogenetically blunting DA and 5HT reward responses simultaneously blocked learning about a reward-predictive cue. Optogenetically reproducing both DA and 5HT responses to reward, but not either one alone, was sufficient to drive the acquisition of new associations. Altogether, these results demonstrate that striatal integration of inverse DA and 5HT signals is a crucial mechanism gating associative learning.

---

Survival depends on an animal’s ability to seek rewards and learn about environmental cues that predict them. From insects to primates, the neuromodulators DA and 5HT have been found to play key roles in this Pavlovian (i.e. cue-outcome) learning process by signaling the presence of rewards (unconditioned stimuli; US) or reward-predictive cues (conditioned stimuli; CS) and regulating synaptic plasticity mechanisms thought to underlie formation of CS-US associations^1, 6, 7, 12, 14–16^. In mammals, DA and 5HT neurons’ shared connectivity with limbic structures^17–19^ suggests that these circuits are coordinately modulated during learning, but precisely how DA and 5HT interactions contribute to the formation of new associations remains a matter of debate.

Historically, this debate has centered around two primary ideas. The synergy hypothesis posits that DA and 5HT signals carry information about reward expectation on short and long timescales, respectively^10, 12, 20^, thereby synthesizing theories about DA’s function in temporal difference learning^21, 22^ and 5HT’s role in regulating mood^23^. The opponency hypothesis proposes that DA invigorates and 5HT suppresses behavioral activation^11, 13, 20, 24–26^ to optimize reward seeking with respect to intertemporal choice tradeoffs^27–29^ or under the threat of punishment^25, 30, 31^, such that imbalances between these processes could lead to compulsion and addiction^32^. So far, however, efforts to test ideas about DA and 5HT interactions directly have been stymied by the difficulty of precisely manipulating multiple neuromodulatory systems at the same time. Here, we present a double transgenic strategy enabling genetic access to the brain’s DA and 5HT systems in a single animal and leverage this advance to reveal that the striatum integrates opponent ventral tegmental area DA (VTA^DA^) and dorsal raphe 5HT (DR^5HT^) signals that coordinately control reinforcement during appetitive learning.

## Simultaneous access to DA and 5HT circuits

We established a mouse model enabling orthogonal genetic access to DA and 5HT neurons by crossing the DAT-Cre and SERT-Flp mouse lines to produce DAT-Cre^+/-^ / SERT-Flp^+/-^ progeny (**Fig. 1a**). To evaluate the cell-type specificity of this mouse line, we injected DAT-Cre^+/-^ / SERT-Flp^+/-^ mice with viral vectors encoding Cre-dependent mCherry in the VTA and Flp-dependent EYFP in the DR (**Fig. 1b, c**), and analyzed the colocalization between these fluorophores and immunostains for DA and 5HT cell markers. We observed >90% colocalization between mCherry and tyrosine hydroxylase (TH) in the VTA, and between EYFP and tryptophan hydroxylase 2 (TpH) in the DR (**Fig. 1d-i**). Importantly, control mice that received the same viruses but into the opposite target regions showed negligible EYFP expression in the VTA and lacked mCherry expression in DR^5HT^ neurons (**Extended Data Fig 1a-d**). Together, these data confirmed that DAT-Cre^+/-^ / SERT-Flp^+/-^ mice enable simultaneous, independent genetic access to midbrain DA and 5HT systems.

**Fig. 1:**
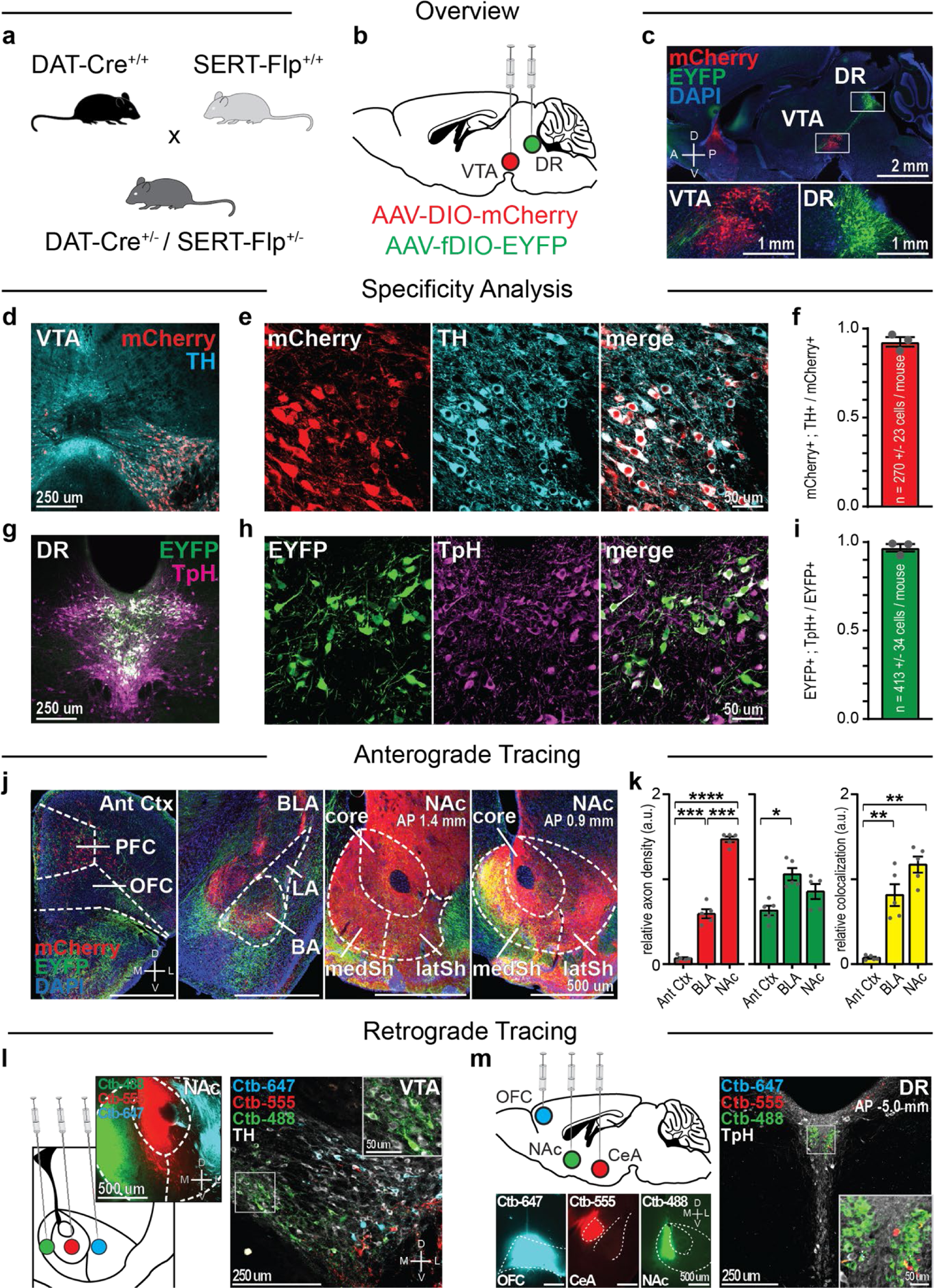
Mapping convergent VTA^DA^ and DR^5HT^ inputs to the NAc. **a**, Schematic describing generation of DAT-Cre^+/-^ / SERT-Flp^+/-^ mice enabling independent genetic access to DA and 5HT neurons. **b**, Viral strategy for labeling VTA^DA^ neurons with mCherry and DR^5H^^T^ neurons with EYFP. **c**, Sagittal section from an example mouse depicting mCherry-expressing neurons in the VTA and EYFP-expressing neurons in the DR. **d-e**, Example images showing colocalization between Cre-dependent mCherry and TH in the VTA. **f**, Cell-type specificity quantification for VTA^DA^ neurons in the DAT-Cre^+/-^ / SERT-Flp^+/-^ line (n=3 mice). **g-h**, Example images showing colocalization between Flp-dependent EYFP and TpH in the DR. **i**, Cell-type specificity quantification for DR^5HT^ neurons in the DAT-Cre^+/-^ / SERT-Flp^+/-^ line (n=3 mice). **j**, Example images of VTA^DA^ and DR^5HT^ inputs to limbic structures. **k**, Density of DA axons (left), 5HT axons (center), and colocalization between DA and 5HT axons (right) across limbic structures (DA axons, one-way RM ANOVA: F_(1.090, 4.359)_=268.4, P< 0.0001; Holm-Sidak tests: Ant Ctx vs NAc, P<0.0001; Ant Ctx vs BLA, P=0.0009; NAc vs BLA, P=0.0009; 5HT axons, one-way RM ANOVA: F_(1.354, 5.415)_=5.965, P=0.0494; tests: Ant Ctx vs BLA, P=0.0370; all other comparisons, P>0.05; colocalization, one-way RM ANOVA: F_(1.014, 4.056)_=24.67, P=0.0073; Holm-Sidak tests: Ant Ctx vs NAc, P=0.0012; Ant Ctx vs BLA, P=0.0082; all other comparisons, P>0.05; n=5 mice). **l**, Left, injection strategy to label projection-defined DA subsystems. Right, example image showing retrogradely labeled VTA^DA^ neurons. **m**, Left, injection strategy to label projection-defined 5HT subsystems. Right, example image showing retrogradely labeled DR^5HT^ neurons. Note the lack of colocalization between Ctb-488 and the other two tracers in **l-m**. Throughout, data are shown as mean +/- s.e.m. and significance is shown as *P<0.05, **P<0.01, ***P<0.001, ****P<0.0001.

## Identifying sites of DA and 5HT convergence

Next, we performed anterograde axon tracing to identify hotspots of VTA^DA^ and DR^5HT^ input convergence in limbic structures important for associative learning. Labeling VTA^DA^ neurons with mCherry and DR^5HT^ neurons with EYFP revealed partially overlapping axon fields in the anterior cortex (Ant Ctx), basolateral amygdala (BLA), and nucleus accumbens (NAc) (**Fig. 1j**). We found variable amounts of colocalization between VTA^DA^ and DR^5HT^ axons across these limbic structures and their respective subregions, with the densest overlap observed in the posterior medial shell of the NAc (NAc^pmSh^) (**Fig. 1k** and **Extended Data Fig. 1e-h**). Retrograde tracing with fluorescently tagged cholera toxin confirmed that DA inputs to the NAc^pmSh^ arise from medial VTA^DA^ neurons that are distinct from NAc core-(NAc^core^) and lateral shell-(NAc^latSh^) projecting subpopulations^18, 33^ (**Fig. 1l**). Surprisingly, we found that 5HT inputs to the NAc also originate from a unique group of DR^5HT^ neurons that was distinct from previously described DR^5HT^ subsystems^34^ and tightly concentrated in the dorsomedial, posterior DR^35^ (**Fig. 1m** and **Extended Data Fig. 1i-r**). Based on these data we focused on the NAc^pmSh^ as the region best positioned to integrate convergent DA and 5HT signals during reward learning.

## DA and 5HT input dynamics during learning

With an appropriate target brain region identified, we then asked how striatal VTA^DA^ and DR^5HT^ inputs respond during a reward learning task. Using DAT-Cre^+/-^ / SERT-Flp^+/-^ mice, we expressed the red calcium indicator RCaMP2 in VTA^DA^ neurons and the green calcium indicator GCaMP6m in DR^5HT^ neurons. This allowed us to simultaneously record the activity of VTA^DA^ and DR^5HT^ axons in the NAc^pmSh^ as mice learned a new cue-reward association (**Fig. 2a-c** and **Extended Data Fig. 2a**). In this task, the CS (a compound sound and port-light cue) predicted delivery of sucrose (US) into the reward port. Mice successfully acquired the CS-US association as evidenced by a reduced latency to collect the reward and an increase in the number of port entries following CS-onset late in training. Aligning the photometry traces to CS-onset and reward consumption showed that VTA^DA^ neurons projecting to the NAc^pmSh^ did not acquire responses to a reward-predictive cue but were excited by rewards, in agreement with a previous report of DA activity in this striatal subregion^36^ (**Fig. 2d, e** and **Extended Data Fig. 2b**). 5HT inputs to the NAc also did not respond to the CS but were inhibited by rewards, consistent with the finding that these inputs arise from DR^5HT^ neurons distinct from previously described subpopulations known to be excited by appetitive stimuli^34^ (**Fig. 2f, g** and **Extended Data Fig. 2c**). Thus, we find dramatically different, inverse response profiles between DA and 5HT inputs to the NAc^pmSh^ (**Extended Data Fig. 2d, e**).

**Fig. 2:**
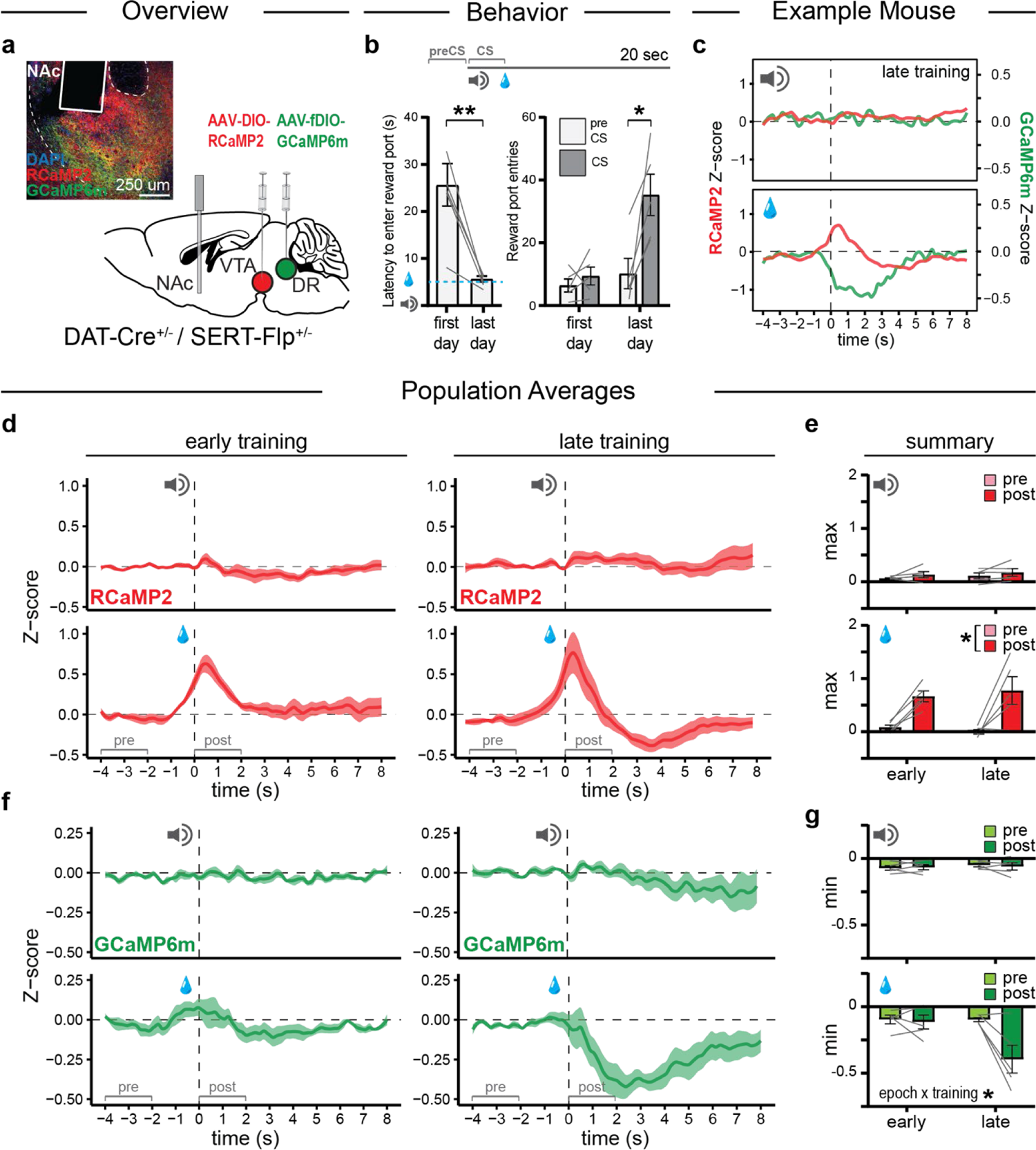
Convergent DA and 5HT inputs to NAc show inverse responses to rewards. **a**, Surgical strategy to record VTA^DA^ and DR^5HT^ axon activity in the NAc. Inset, image of the recording site from a representative mouse. **b**, Top, schematic of the conditioning task. CS-(compound sound and light cue) onset predicted US (sucrose solution) delivery into a reward port. Mice acquired the CS-US association as evidenced by a shortened latency to collect rewards (left, paired t-test: t_(4)_=4.828, P=0.0085, n=5 mice) and an increased number of port entries following CS-onset late in training (right, two-way RM ANOVA: training x period interaction, F_(1.000, 4.000)_=8.053, P=0.0470, n=5 mice; Holm-Sidak tests, preCS vs CS: first day, P=0.4263; last day, P=0.0499). **c**, Simultaneously recorded DA and 5HT responses from an example mouse. **d-g**, Population recordings and peak Z-score quantifications of VTA^DA^ (**d,e**) and DR^5HT^ (**f,g**) activity aligned to CS onset (top) or reward consumption (bottom) during early (left) and late (right) training. VTA^DA^ axons did not respond to the CS (top, two-way RM ANOVA: main effect of training, F_(1.000,4.000)_=0.8768, P=0.4021; main effect of period, F_(1.000,4.000)_=1.852, P=0.2451; training x period interaction, F_(1.000,4.000)_=0.0592, P=0.8197; n=5 mice) and were excited by rewards (bottom, two-way RM ANOVA: main effect of training, F_(1.000,4.000)_=0.0356, P=0.8596; main effect of period, F_(1.000,4.000)_=19.32, P=0.0117; training x period interaction, F_(1.000,4.000)_=0.6918, P=0.4523; n=5 mice). DR^5HT^ axons did not respond to the CS (top, two-way RM ANOVA: main effect of training, F_(1.000,4.000)_=0.3209, P=0.6013; main effect of period, F_(1.000,4.000)_=0.0446, P=0.8430; training x period interaction, F_(1.000,4.000)_=0.2335, P=0.6542; n=5 mice) and were inhibited by rewards (bottom, two-way RM ANOVA: training x period interaction, F_(1.000,4.000)_=8.997, P=0.0400, n=5 mice; Holm-Sidak tests, pre vs post: early, P=0.7303; late, P=0.0640).

## Inverse DA and 5HT responses gate reinforcement

The previous experiments assessing DA and 5HT input activities suggest that the two modulatory systems encode rewards in opposite manners. However, these passive observations of input activity are insufficient to conclude that these dramatic differences in DA and 5HT reward signals have important behavioral significance. To directly test the necessity of VTA^DA^ and DR^5HT^ reward responses for reward learning, we performed a loss-of-function experiment designed to blunt either DA or 5HT signals alone or both together in response to an appetitive US. Specifically, we used DAT-Cre^+/-^ / SERT-Flp^+/-^ mice to express halorhodopsin (NpHR) or an EYFP control in VTA^DA^ neurons and channelrhodopsin (ChR2) or an EYFP control in DR^5HT^ neurons in a two-by-two design and implanted them with bilateral optical fibers in the NAc^pmSh^. This enabled us to use long pulses of red light to blunt VTA^DA^ excitatory reward responses and short pulses of blue light to blunt DR^5HT^ inhibitory reward responses in the NAc^pmSh^ during reward consumption in the sucrose conditioning task (**Fig. 3a**). Over the course of training, mice in the single opsin groups – in which either DA inputs were inhibited (EYFP/NpHR) or 5HT inputs were excited (ChR2/EYFP) – obtained an equivalent number of rewards and collected them just as efficiently as the control (EYFP/EYFP) group (**Fig. 3b-d).** In contrast, mice in the double opsin (NpHR/ChR2) group – in which both optogenetic manipulations occurred simultaneously – obtained fewer rewards, took longer to collect them, and made fewer port entries (**Fig. 3b-d**).

**Fig. 3:**
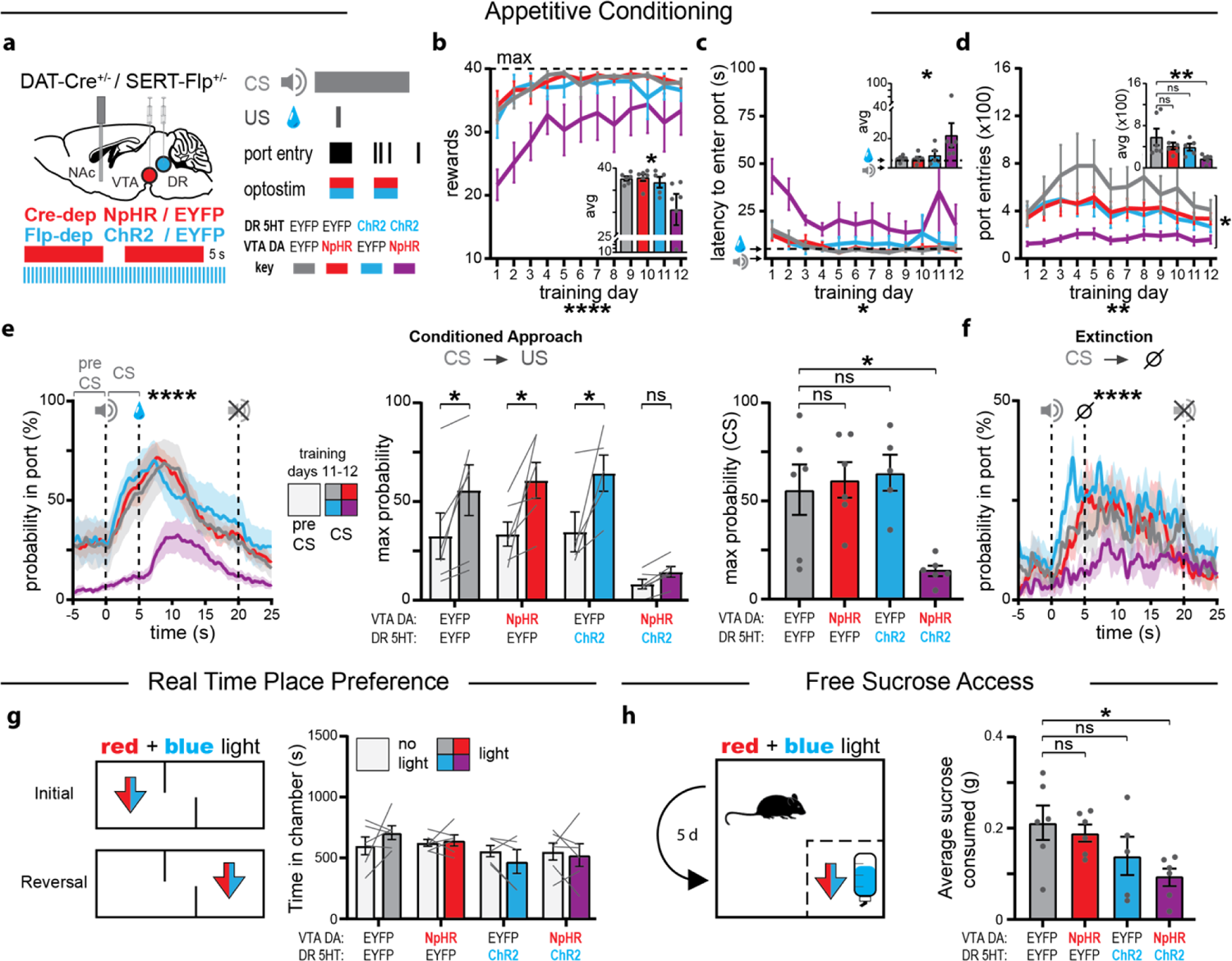
Coordinated blunting of DA and 5HT reward responses disrupts learning. **a**, Optogenetic strategy to blunt VTA^DA^ and/or DR^5HT^ reward responses during learning. **b**, Rewards as a function of training (two-way RM ANOVA: main effect of virus, F_(3,19)_=3.112, P=0.0507; main effect of time, F_(2.247,42.68)_=10.94, P<0.0001, virus x time interaction, F_(33,209)_=1.399, P=0.0840). Inset, average rewards across days (Kruskal-Wallis test: P=0.0326; Dunn’s test: EYFP/EYFP vs NpHR/EYFP, P>0.9999; EYFP/EYFP vs EYFP/ChR2, P>0.9999; EYFP/EYFP vs NpHR/ChR2, P=0.0896). **c**, Latency to collect rewards as a function of training (two-way RM ANOVA: main effect of virus, F_(3,19)_=2.920, P=0.0606; main effect of time, F_(1.678,31.88)_=3.543, P=0.0481, virus x time interaction, F_(33,209)_=0.9067, P=0.6178). Inset, average latency across days (Kruskal-Wallis test: P=0.0317; Dunn’s test: EYFP/EYFP vs NpHR/EYFP, P>0.9999; EYFP/EYFP vs EYFP/ChR2, P>0.9999; EYFP/EYFP vs NpHR/ChR2, P=0.0514). **d**, Port entries as a function of training (two-way RM ANOVA: main effect of virus, F_(3,19)_=3.314, P=0.0421; main effect of time, F_(3.249,61.73)_=4.382, P=0.0061; virus x time interaction, F_(33,209)_=0.8065, P=0.7654). Inset, average port entries across days (Kruskal-Wallis test: P=0.0091; Dunn’s test: EYFP/EYFP vs NpHR/EYFP, P>0.9999; EYFP/EYFP vs EYFP/ChR2, P>0.9999; EYFP/EYFP vs NpHR/ChR2, P=0.0087). **e**, Probability of occupying the reward port as a function of time (left, Mixed-effects model: virus x time interaction, χ^2^ =1273.6, P<0.0001). Peak probability as a function of CS-onset (center, two-way RM ANOVA: main effect of virus, F_(3,19)_=5.496, P=0.0069; main effect of epoch, F_(1,19)_=26.84, P<0.0001; virus x epoch interaction, F_(3,19)_=1.658, P=0.2097; Holm-Sidak tests, preCS vs CS: EYFP/EYFP, P=.0200; NpHR/EYFP, P=0.0134; EYFP/ChR2, P=0.0134; NpHR/ChR2, P=0.4609) and following CS-onset (right, Kruskal-Wallis test: P=0.0092; Holm-Sidak tests: EYFP/EYFP vs NpHR/EYFP, P>0.9999; EYFP/EYFP vs EYFP/ChR2, P>0.9999; EYFP/EYFP vs NpHR/ChR2, P=0.0361). **f**, Same as **e** (left), but during extinction (Mixed-effects model: virus x time interaction, χ^2^ =389.05, P<0.0001). **g-h**, Loss-of-function manipulations had no inherent valence (**g**, two-way RM ANOVA: main effect of virus, F_(3,19)_=1.498, P=0.2472; main effect of light, F_(1,19)_=0.0060, P=0.9389; virus x light interaction, F_(3,19)_=1.194, P=0.3386) but reduced sucrose consumption (**h**, one-way ANOVA: F_(3,19)_=3.262, P=0.0442; Holm-Sidak tests: EYFP/EYFP vs NpHR/EYFP, P=0.5915; EYFP/EYFP vs EYFP/ChR2, P=0.2118; EYFP/EYFP vs NpHR/ChR2, P=0.0290). All panels, n=5-6 mice/group.

Late in training, the control and single opsin groups all showed strong evidence of having acquired the CS-US association as evidenced by an increased probability of occupying the reward port following CS-onset (**Fig. 3e, f**). By contrast, mice in the double opsin group did not adapt their reward seeking behavior in response to CS-onset and were least likely to occupy the reward port during the anticipatory period between CS-onset and reward delivery; an effect that persisted even in an extinction session without optogenetic manipulations (**Fig. 3e, f**). The profound disruption of learning that occurred when VTA^DA^ and DR^5HT^ reward responses were blunted simultaneously, but not individually, suggests that the coordinated changes in DA and 5HT input activity are required for optimal learning of the CS-US association.

Notably, this effect of the combined optogenetic manipulations cannot be explained by affective (i.e. valence) or locomotor changes driven by the optogenetic manipulations themselves, since in the same subjects these manipulations had no effects in real time place preference (RTPP) or open field test control assays (**Fig. 3g**, **Extended Data Fig. 3a**).

Furthermore, there were no differences between groups in the extent of the modest food restriction we employed (**Extended Data Fig. 3b**) and all optical fiber placements were confirmed to have been targeted correctly (**Extended Data Fig. 3c-j**). Instead, additional experiments in which mice were given free access to sucrose rewards suggest that the learning deficit observed in the Pavlovian conditioning task may be driven by a reduction in the hedonic or reinforcing properties of the US in mice with blunting of both endogenous VTA^DA^ and DR^5HT^ reward responses (**Fig. 3h**).

## Integration of DA and 5HT signals drives learning

Next, we asked whether VTA^DA^ and/or DR^5HT^ reward signals in the NAc^pmSh^ are sufficient to drive associative learning. To reproduce each neuromodulatory response to reward alone or both together, we expressed ChR2 or EYFP in VTA^DA^ and NpHR or EYFP in DR^5HT^ neurons of DAT-Cre^+/-^ / SERT-Flp^+/-^ mice in a two-by-two design and implanted them with bilateral optical fibers in the NAc^pmSh^. We tested these mice on an optogenetic conditioned place preference (CPP) assay where one side of a two-chambered box was paired with optostimulation/inhibition and the other side was paired with no optogenetic manipulation (**Fig. 4a**). Neither DR^5HT^ inhibition nor VTA^DA^ stimulation in the NAc^pmSh^ were individually sufficient to drive CPP in the single opsin groups (EYFP/NpHR, ChR2/EYFP), whereas both manipulations delivered together produced a place preference in every mouse that expressed both opsins (ChR2/EYFP) (**Fig. 4b, c**). This result suggests that the integration of VTA^DA^ and DR^5HT^ reward responses in the NAc^pmSh^ is important for animals to learn a new association.

**Fig. 4:**
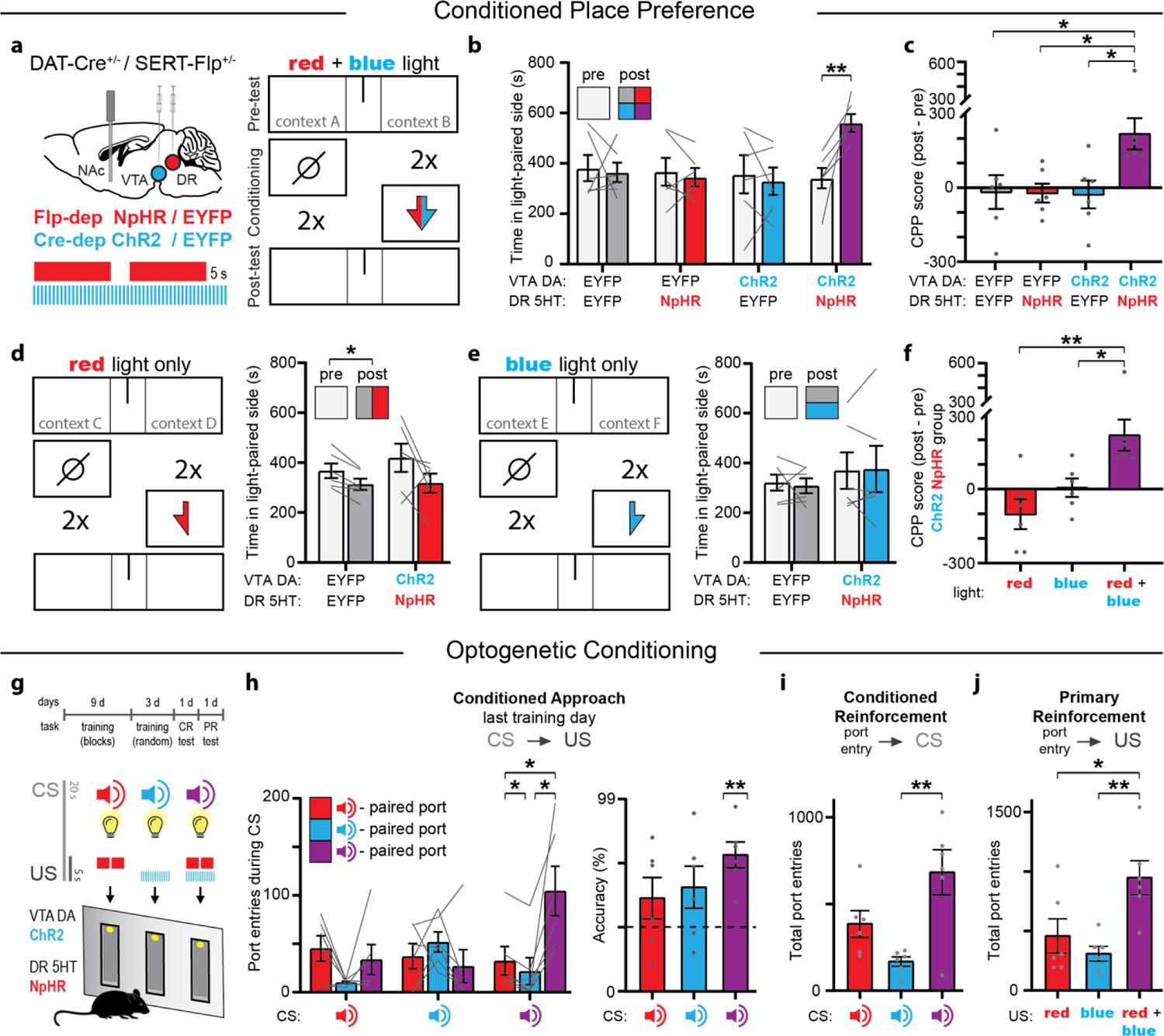
Integration of convergent DA and 5HT reward responses drives learning. **a**, Left, optogenetic strategy to reproduce VTA^DA^ and/or DR^5HT^ reward responses. Right, CPP paradigm schematic. **b-c**, VTA^DA^ excitation and DR^5H^^T^ inhibition together, but not either manipulation alone, produced CPP (**b**, two-way RM ANOVA: virus x light interaction, F_(3,20)_=4.347, P=0.0164; Holm-Sidak tests, pre vs post: EYFP/EYFP, P=0.9550; EYFP/NpHR, P=0.9550, ChR2/EYFP, P=0.9550; ChR2/NpHR, P=0.0046; **c**, one-way ANOVA: F_(3,20)_=4.347,P=0.0164; Holm-Sidak tests: EYFP/EYFP vs ChR2/NpHR, P=0.0407; EYFP/NpHR vs ChR2/NpHR, P=0.0407; ChR2/EYFP vs ChR2/NpHR, P=0.0407; all other comparisons P>0.05). **d-f**, DR^5HT^ inhibition or VTA^DA^ excitation alone did not produce CPP in the same mice that previously showed CPP in response to both manipulations together (**d**, two-way RM ANOVA: main effect of virus, F_(1,10)_=0.3929, P=0.5448; main effect of light F_(1,10)_=6.267, P=0.0313; virus x light interaction, F_(1,10)_=0.5688, P=0.4681; **e**, two-way RM ANOVA: main effect of virus, F_(1,10)_=0.4574, P=0.5142; main effect of light F_(1,10)_=0.0197, P=0.8913; virus x light interaction, F_(1,10)_=0.1408, P=0.7154; **f**, one-way RM ANOVA: F_(2,15)_=8.742, P=0.0030; Holm-Sidak tests: red vs red+blue, P=0.0028; blue vs red+blue, P=0.0308; all other comparisons P>0.05). Purple bars in **c** and **f** represent the same data. **g**, Optogenetic conditioning paradigm with three CS-(compound sound and port light cues) US-(VTA^DA^ stimulation and/or DR^5HT^ inhibition) pairs. **h-i**, Conditioned approach behavior evoked by the CS paired with both manipulations was stronger (**h**, left, two-way RM ANOVA: CS x port interaction, F_(1.878,9.391)_=12.64, P=0.0024; Holm-Sidak tests, CS-red+blue: red port vs blue port, P=0.0420; red port vs red+blue port, P=0.0420; blue port vs red+blue port, P=0.0420; all other comparisons P>0.05), and more accurate (right, Bonferroni-corrected, one-sample t-tests with hypothetical mean 33.33: CS-red, P=0.6867, CS-blue, P=0.3585, CS-red+blue, P=0.0081) than that evoked by CSs paired with either manipulation alone. **i**, The CS paired with both manipulations became a potent conditioned reinforcer (Friedman’s test, P=0.0012; Dunn’s test: CS-blue vs CS-red+blue, P=0.0078; all other comparisons P>0.05). **j**, Mice preferred VTA^DA^ stimulation and DR^5HT^ inhibition delivered together to either manipulation alone (Friedman’s test, P=0.0055; Dunn’s test: CS-blue vs CS-red+blue, P=0.0078; CS-red vs CS-red+blue, P=0.0418; all other comparisons P>0.05). All panels, n=6 mice/group.

To further test that both VTA^DA^ and DR^5HT^ reward signals must be present in the NAc^pmSh^ to drive CPP, we performed several additional experiments. First, we again characterized the affective and locomotor effects of the optogenetic manipulations themselves. In the RTPP test, we found that VTA^DA^ stimulation in the NAc^pmSh^ was only rewarding when it was delivered together with DR^5HT^ inhibition, suggesting both signals must be present to drive reinforcement (**Extended Data Fig. 4a**). Both the ChR2/EYFP and ChR2/NpHR groups showed a locomotor effect in the open field test (**Extended Data Fig. 4b**); results that confirm that manipulations to optogenetically stimulate VTA^DA^ inputs in the NAc^pmSh^ were functional. Second, in the same mice that had previously shown a robust CPP in response to both manipulations together (**Fig. 4b**), we asked whether VTA^DA^ stimulation and DR^5HT^ inhibition alone could drive CPP. Specifically, we took the mice in the ChR2/NpHR group and repeated the CPP assay twice with only blue and only red light, using distinct contextual cues in each iteration of the test to minimize the chances of carryover learning. Neither VTA^DA^ input stimulation alone nor DR^5HT^ input inhibition alone generated CPP on their own (**Fig. 4d, e**), even in the same mice that had previously learned a strong preference for both manipulations delivered simultaneously (**Fig. 4f**).

Finally, we extended these findings to more complex forms of associative learning using an optogenetic pavlovian conditioning task^3^. We placed ChR2/NpHR mice from the double opsin group into an operant chamber with three ports, each equipped with its own port light. Mice were then presented with three cue-outcome pairings where compound sound and port-light CSs predicted delivery of optogenetic USs consisting of VTA^DA^ stimulation alone, DR^5HT^ inhibition alone, or both together (**Fig. 4g**). Crucially, US delivery was not contingent upon port entry, allowing us to use port entries as a measure of conditioned approach; a behavioral read-out of learning. Despite being trained on an equivalent number of trials of each type (**Extended Data Fig. 4c**), mice acquired conditioned approach responses that were stronger and more accurate in response to the CS that predicted VTA^DA^ stimulation and DR^5HT^ inhibition together compared to the CSs paired with either manipulation alone (**Fig. 4h**). Furthermore, the CS paired with VTA^DA^ stimulation and DR^5HT^ inhibition together became a more potent conditioned reinforcer than the CS paired with either manipulation alone (**Fig. 4i**). As a final direct test of the relative potency of VTA^DA^ stimulation alone, DR^5HT^ inhibition alone, or both together as primary reinforcers, mice were given the choice to nose poke to receive any one of these three manipulations and strongly preferred to receive the two optogenetic manipulations together compared to either one alone (**Fig. 4j**). After confirming the accuracy of our optical fiber placements (**Extended Data Fig. 4d-k**), we conclude that inverse VTA^DA^ and DR^5HT^ signals converging onto the NAc^pmSh^ are integrated to drive learning.

## Discussion

Together our data suggest that optimal associative learning requires inverse VTA^DA^ and DR^5HT^ signals converging onto the NAc^pmSh^. How do these findings fit with proposed models of DA and 5HT interactions? On the whole, our data are incongruent with the synergy across timescales theory^10, 12, 21^ given that the DA and 5HT reward responses we recorded occurred with similar timescales on the order of seconds. Furthermore, while these phasic responses do not necessarily preclude the existence of additional tonic signals carrying reward information on longer timescales (which could be difficult to detect in our axon calcium activity recordings), the results of our optogenetic manipulations strongly suggest this is unlikely. Neither stimulation nor inhibition of VTA^DA^ or DR^5HT^ NAc^pmSh^ inputs alone was inherently rewarding or aversive, as would be expected if either one of these modulators were signaling long-term reward value or expectation.

Instead, our results demonstrating an inverse relationship between DA and 5HT reward signals support models of DA and 5HT opponency^11, 13, 21, 24–26^. For example, our optogenetic manipulation experiments suggest that opposing VTA^DA^ and DR^5HT^ responses must coincide in the NAc^pmSh^ to drive learning about reward-predictive cues in CPP and pavlovian conditioning assays. This effect is unlikely to be explained by coordinated DAergic and 5HTergic control of behavioral activation and inhibition as proposed by earlier theories^11, 13, 20, 24–26^, given that manipulations of DR^5HT^ inputs alone did not affect locomotion. Rather, integration of VTA^DA^ and DR^5HT^ NAc^pmSh^ inputs was sufficient to drive real-time place preference and coordinately blunting both modulatory reward responses reduced free reward consumption without being inherently aversive. Together these results suggest that VTA^DA^ and DR^5HT^ inputs to NAc^pmSh^provide coordinated, opponent modulatory control over reinforcement.

We intentionally performed our manipulations in the NAc^pmSh^, the target region in which we found the most anatomical overlap between VTA^DA^ and DR^5HT^ axons. This projection-specific strategy resulted in optogenetic manipulations that did not affect other modulatory circuits engaged by natural rewards, which could explain why learning was spared in the single opsin groups with loss-of-function targeted to either VTA^DA^ or DR^5HT^ reward responses alone.

A potential limitation of this approach is that our conclusions may be restricted to interactions between striatal 5HT inputs and DA inputs arising from noncanonical VTA neurons projecting to the NAc^pmSh^, which are excited by both rewards and punishments but lack responses to reward-predictive cues^36–40^. This may explain why neither stimulation nor inhibition of these specific VTA^DA^ inputs alone was inherently rewarding or aversive. However, recent work showing that inhibition of 5HT release in the dorsal striatum potentiates the reinforcing properties of a cocaine reward^32^ suggests that the opponent relationship we describe between DA and 5HT inputs to NAc^pmSh^is conserved across striatal subregions.

Our experiments directly address a longstanding controversy about the nature of DA and 5HT interactions in a manner not previously possible and reveal a fundamental biological process underlying associative learning. Our results support an updated opponency model where the integration of inverse DA and 5HT signals in the striatum governs reward conditioning by gating reinforcement. We hope this work will motivate further investigations into how the combinatorial actions of DA and 5HT elsewhere in the brain shape other motivated behaviors thought to be under multiplexed neuromodulatory control, including aversive learning^5, 14^ and sociability^41, 42^, with the potential to deepen our understanding of psychiatric diseases characterized by dysfunction in these behaviors.

## METHODS

### Mice

Adult (>6 week old) wild-type C57BL6/J (Jackson Laboratory #000664) and DAT-Cre^+/-^ / SERT-Flp^+/-^ (in-house cross between Jackson Laboratory lines #006660 and #034050 on a C57BL6 background) mice of both sexes were used in this study. All mice had *ad-libitum* access to food and water unless otherwise specified, and all experimental procedures were approved by the Stanford University Administrative Panel on Laboratory Animal Care and the Administrative Panel on Biosafety.

### Stereotactic surgeries

Mice were anesthetized with either isofluorane (4-5% induction, 1-2% maintenance) or a ketamine-(60 mg/kg) dexmedetomidine (0.6 mg/kg) cocktail. Within each experiment, the type of anesthesia used was the same for every mouse. A stereotaxic frame (Kopf instruments) was used to target manipulations to the following structures (coordinates are in mm, with AP and ML relative to bregma and DV relative to the skull surface): DR, AP - 4.6, ML 0, DV - 3.3; VTA, AP - 3.4, ML +/- 0.2, DV - 4.2; NAc medSh, AP +1.0, ML +/-0.7, DV - 4.3 to −4.5; NAc core, AP +1.1, ML 1.2, DV −4.5; NAc latSh, AP +1.1, ML 1.8, DV −4.5; OFC, AP +2.6, ML 1.1, DV −2.5; CeA, AP - 1.5, ML 2.5, DV - 4.5. Viral vectors (∼10^13^ gc/mL for photometry experiments, ∼10^12^ gc/mL otherwise; from the Stanford Gene Vector and Virus core unless otherwise specified) and Ctb retrograde tracers (1 ug/ul; Invitrogen #C34777, #C34776, #C34775) were infused with a syringe pump (Harvard Apparatus) at a rate of 100-200 nL/min and allowed to diffuse for at least 5 min. Incubation times were >5 weeks for viruses and 1-2 weeks for retrograde tracers. Optical fibers (Doric Lenses, 200-400 um diameter, 0.48-0.66 NA) were secured to the skull using screws (Antrin Miniature Specialties) and dental cement (Geristore), and fiber placements were histologically verified post-hoc. Mice with mistargeted fibers, defined as fiber tips outside the NAc medSh or outside the anteroposterior range of 0.8-1.2 mm from bregma where DR^5HT^ axons are dense in the NAc, were excluded from further analysis.

### Histology and Immunohistochemistry

Histological procedures were performed as previously described^1^. Mice were transcardially perfused with 4% (w/v) paraformaldehyde (PFA) in phosphate-buffered saline (PBS) and postfixed in PFA overnight. Coronal sections 50 um thick were immunostained for tyrosine hydroxylase (primary: mouse anti-TH, Millipore #MAB318; secondary: goat anti-mouse 647, Invitrogen #A-21236 or goat anti-mouse 405, Invitrogen #A-31553), tryptophan hydroxylase 2 (primary: rabbit anti-TpH, Novus #NB100-74555; secondary: goat anti-rabbit 546, Invitrogen #A-11035 or goat-anti rabbit 405, Invitrogen #A-31556), EYFP (primary: chicken anti-GFP, Aves #GFP-1020; secondary: goat anti-chicken 488, Invitrogen #A-11039), and/or mCherry (primary: rat anti-mCherry, Invitrogen #M11217; secondary: goat anti-rat 594, Invitrogen #11007). Primary antibodies were used at a concentration of 1:1000 and incubated on a shaker overnight at room temperature. Secondary antibodies were used at a concentration of 1:750 and incubated on a shaker for 2 hours at room temperature.

### Cell-type specificity analysis

DAT-Cre^+/-^ / SERT-Flp^+/-^ mice were injected with 500 nL of AAV-DJ-EF1a-DIO-mCherry unilaterally into the VTA and 500 nL of AAV-DJ-EF1a-fDIO-EYFP into the DR. Cell-type specificity analysis was then performed as previously described^1^. Briefly, coronal sections of the VTA were stained for TH and DR sections were stained for TpH. For every other section, 40x images (∼300 um x 300 um, 40x objective, Nikon A1 confocal microscope) were acquired in each of the dorsomedial, ventromedial, and lateral (unilateral for VTA, bilateral for DR) subregions of the DR or VTA. Cell-type specificity for VTA^DA^ and DR^5HT^ neurons was defined as the fraction of mCherry+ cells that were TH+ and the fraction of EYFP+ cells that were TpH+, respectively. Mice in the orthogonality control group were prepared identically to those in the cell-type specificity experiment but with the targets for the viruses swapped, and alternating DR sections were stained for TH and TpH to confirm that Cre-dependent mCherry expression restricted to DR^DA^ neurons and not present in DR^5HT^ neurons.

### Anterograde anatomical tracing

DAT-Cre^+/-^ / SERT-Flp^+/-^ mice were injected with 400 nL of AAV-DJ-EF1a-DIO-mCherry bilaterally into the VTA and 800 nL of AAV-DJ-EF1a-fDIO-EYFP into the DR. Coronal sections were immunostained for EYFP and mCherry to amplify the signal from labeled axons. Epifluorescence images of the Ant Ctx, anterior BLA (AP - 0.8 mm to −1.5 mm), posterior BLA (AP - 1.5 mm to −2.2 mm), anterior NAc (AP 1.2 mm to 1.6 mm), and posterior NAc (AP 0.8 mm to 1.2 mm; 6-8 samples/region/mouse) were acquired on a Keyence BZ-X800 microscope using a 10x objective and identical imaging settings for all samples. Image analysis was done with ImageJ. Each image was background subtracted twice (first with a rolling ball radius of 50 px, then 25 px) and the green and red channels were binarized using Otsu’s thresholding method^2^. To measure colocalization between VTA^DA^ and DR^5HT^ axons, the binarized green and red images were multiplied together yielding images that only had signal in pixels that were thresholded in both the green and red channels. Since pixels measured ∼2 x 2 um each and both 5HT and DA can signal via volume transmission^3^ over comparable distances^4^, we reasoned that this analysis would reveal regions with cells that are well positioned to receive convergent DA and 5HT inputs. Regions of interest were manually defined based on DAPI signal and the Paxinos mouse brain atlas^5^. For each region, the VTA^DA^ axon density, DR^5HT^ axon density, and axon colocalization were measured as the fraction of segmented pixels in the red, green, and colocalization channels, respectively, and the data for each channel was normalized within mice.

### Retrograde anatomical tracing

Wild-type or DAT-Cre^+/-^ / SERT-Flp^+/-^ mice were injected with 200-300 nL each of Ctb-488, Ctb-555, and Ctb-647. To identify subsystems of VTA^DA^ neurons, retrograde tracers were injected in the NAc medSh, NAc core, and NAc latSh, respectively, and VTA sections were immunostained for TH. To identify subsystems of DR^5HT^ neurons, retrograde tracers were injected in the NAc medSh, CeA, and OFC, respectively, and DR sections were immunostained for TpH. For each DR section, the anteroposterior coordinate was estimated using the Paxinos mouse brain atlas^5^, 40x images (∼300 um x 300 um, 40x objective, Nikon A1 confocal microscope) were acquired in each of the dorsomedial, ventromedial, lateral left, and lateral right subregions of the DR, and colocalization between the tracers and TpH was manually quantified. For retrograde tracing control experiments, mice were injected with 200-300 nL of a 1:1:1 cocktail of all three retrograde tracers in the NAc medSh and the DR was imaged and analyzed as above.

### Fiber photometry

DAT-Cre^+/-^ / SERT-Flp^+/-^ mice were injected with 1000 nL of AAV-DJ-EF1a-DIO-RCaMP2 unilaterally into the VTA and 1000 nL of AAV-DJ-EF1a-fDIO-GCaMP6m into the DR and implanted unilaterally with an optical fiber in the NAc medSh (ipsilateral to the VTA injection). Data collection and preprocessing followed procedures described previously^6^ that were modified to enable simultaneous recording in the red and green channels. Briefly, mice were hooked up through low-autofluorescence patch cords (Doric) to a photodetector (Newport #2151). GCaMP and RCaMP sensors were excited by frequency modulated 405 nm, 465 nm, and 560 nm light with LED power settings held constant across all mice. The resulting fluorescence signals were sampled at 1.0173 KHz, band pass filtered, demodulated, and recorded using a signal processor (RZ5P, Tucker-Davis Technologies). GCaMP and RCaMP traces were detrended (by subtracting out the linear fit between them and the 405 nm channel, respectively), digitally filtered, aligned to behavioral events of interest, z-scored, and smoothened using local regression over a sliding window of 100 ms.

### Optogenetic stimulation and inhibition

For loss-of-function experiments, DAT-Cre^+/-^ / SERT-Flp^+/-^ mice were injected with 500 nL of either AAV-DJ-ef1a-DIO-NpHR3.0-eYFP or AAV-DJ-ef1a-DIO-eYFP bilaterally into the VTA and 1000 nL of AAV-DJ-EF1a-fDIO-ChR2 or AAV-DJ-ef1a-fDIO-eYFP into the DR and implanted bilaterally with optical fibers in the NAc medSh. For gain-of-function experiments, DAT-Cre^+/-^ / SERT-Flp^+/-^ mice were injected with 500 nL of either AAV-DJ-ef1a-DIO-ChR2-eYFP or AAV-DJ-ef1a-DIO-eYFP bilaterally into the VTA and 1000 nL of AAV-8-nEF-CoffFon-NpHR3.3-eYFP (Addgene #137154) or AAV-DJ-ef1a-fDIO-eYFP into the DR and implanted bilaterally with optical fibers in the NAc medSh. Prizmatix Pulsers and STSI LEDs were used to deliver blue (450 nm, 11-14 mW, 5 ms pulses at 20 Hz) and/or red (620 nm, 5-8 mW, in a 2s on 0.5 s off pattern unless otherwise specified) light through a single patch cord that mice were connected to via a rotary joint (Doric).

### General behavioral procedures

Mice received at least two sessions of habituation to handling, one session of habituation to each new behavioral arena, and an additional ∼2 min of habituation time once in the behavioral apparatus prior to each behavioral testing session. In experiments involving optical fibers, mice were also habituated to being tethered. In experiments involving sucrose rewards, mice were pre-exposed to sucrose solution, food restricted to ∼85% of their *ad-libitum* body weight prior to beginning and for the duration of the behavioral task, and fed daily after the completion of behavioral testing. Blinding was not used because all behavioral measurements were made in an automated way without any manual scoring. Sucrose conditioning and optogenetic conditioning tasks were run using Med-PC IV/V software; all other behavioral tasks were run using the Biobserve behavioral tracking program.

### Appetitive conditioning with sucrose

Sucrose conditioning took place inside of sound insulated operant chambers equipped with a red house light, sound generating speakers, and a reward port fed by a syringe pump (all from Med Associates). The CS consisted of a compound port light and sound cue (fiber photometry experiments: white noise or a 4500 Hz tone, counterbalanced; loss-of-function experiments: white noise only; all 70-80 dB) that lasted for 20 s. The US was ∼12 uL of sucrose solution (32% w/v) delivered into the reward port 5 s after CS-onset. Mice needed to pick up the previous reward before the next trial could begin. The ITI was variable with an average of 40 s, for a maximum of 40 rewards per 40 min session.

For fiber photometry experiments, mice received 5 days of initial training and then continued to receive daily training sessions until they attained 30/40 possible rewards for 3 consecutive days (minimum 8 training days, maximum 14, average ∼10). Early and late training photometry data shown correspond to the first 3 and last 3 days of training for each mouse, respectively. Reward consumption was defined as the first reward port entry following each reward delivery.

For loss-of-function experiments, all mice received 12 days of training, and optostimulation/inhibition was aligned to reward consumption. Specifically, reward port entries made while the CS was on triggered 5 s of optostimulation/inhibition. If no reward port entry was made while the CS was on, then a single 5 s bout of optostimulation/inhibition was delivered upon the next port entry. Data from one training session between days 9 and 10 during which a software crash caused the LEDs not to work on some trials was excluded from analysis. The probability of being in the reward port as a function of time was calculated as the fraction of trials during which each mouse was occupying the reward port at each point in time within a trial, measured in 10 ms bins and smoothened within mice using local regression (span ∼1 s). Extinction sessions had an average ITI of 80 s for a total of 20 trials where neither US or optostimulation/inhibition were delivered.

### Real-time place preference

Real-time place preference tests were done using a 70 cm x 23 cm box divided into three chambers. The center chamber had white walls and smooth floors, while the left and right chambers had distinct visual cues on the walls and floors with different textures. Mice were confined to the center chamber for a ∼2 min habituation period. Then, the barriers were removed to begin the initial phase of the test during which mice were free to explore all three chambers for 15 min and one side chamber (left or right, counterbalanced) was paired with optostimulation/inhibition. Immediately after completion of the initial phase, the side that was paired with optostimulation/inhibition was swapped and mice were free to explore all three chambers for an additional 15 min reversal phase. One mouse that failed to explore all three chambers of the box was excluded from analysis.

### Free sucrose access

Mice were placed into a 40 x 25 cm behavioral arena with a sipper bottle in one corner of the box and given free access to 32% sucrose solution (w/v) for 25 min/day for 5 days. Optogenetic manipulations were delivered for the entire time that each mouse occupied a 10 x 10 cm zone around the sucrose sipper bottle. To ensure maximal inhibition of the VTA^DA^ response to reward, the red light illumination in this experiment was constant as opposed to being delivered in the 2 s on, 0.5 s off pattern used for all other optogenetic experiments. The amount of sucrose solution consumed was measured as the change in weight of the sucrose bottle before and after the task, minus the average change in weight of an identical bottle placed into a box with no mouse in it (to control for the small amount of sucrose that can drip out when the bottle is moved). In 3/115 testing sessions when mice partially chewed through the rubber stopper on the sucrose bottle and spilled some of the sucrose solution inside, the amount consumed was estimated as the change in the mouse’s weight before and after the task. Data shown correspond to the amount of dry sucrose consumed by weight, calculated as the amount of sucrose solution consumed by weight multiplied by the concentration of the solution (32%) and divided by the density (1.0756 g/ml).

### Open field test

Mice were placed into a 40 x 40 cm behavioral arena and allowed to explore freely for 12 mins. Optogenetic manipulations were delivered in four alternating light-on and light-off epochs that lasted 3 mins each and which epoch came first was counterbalanced across mice.

### Conditioned place preference

A 70 cm x 23 cm box was divided into two chambers with different visual cues on the walls and floors with different textures. On the first day of the task, mice underwent a pre-test where they were free to explore the entire box for 15 min. On the second day of the task, mice received two conditioning sessions where first they were confined to one chamber for 25 min and administered optostimulation/inhibition, and then >4 h later they were confined to the other chamber and administered nothing. On the third day, the conditioning procedure was repeated but with the order of the sessions (optostimulation/inhibition or nothing) counterbalanced relative to the first day. On the fourth day the mice underwent a post-test that was identical to the pre-test, and the amount of time spent on the light-paired side before and after conditioning was compared. Which chamber was paired with optostimulation/inhibition was counterbalanced across mice in an unbiased design. For mice that underwent multiple CPP experiments, the visual and textile cues in the boxes were changed for each new experiment.

### Optogenetic conditioning

Optogenetic conditioning experiments were carried out in operant chambers enclosed in sound attenuating boxes and equipped with a sound generator and three reward ports, each with their own reward port light. Mice were presented with three CS-US pairs where CSs were compound sound (white noise, 4500 Hz pure tone, or 40 Hz clicks; all ∼70 dB) and port-light (left, center, or right port) cues and USs consisted of red light stimulation, blue light stimulation, or both. Each CS lasted 20 s, each US lasted 5 s, and the CS and US co-terminated. CS-US pairings were counterbalanced across mice, and US delivery was not contingent upon entry into any port.

Mice underwent 12 training sessions lasting 14 hours each and consisting of ∼420 trials (variable ITI with average 100s) divided approximately evenly between the three trial types. During the first 9 days of training, trials were delivered into blocks of a single trial type (∼90 min/block, 3 blocks of each trial type per session, delivered in randomized order). On the last 3 days of training, trials of different types were interleaved and delivered in randomized order.

Data from one day of training when a software crash caused the optostimulation/inhibition to fail for some mice was excluded from analysis. Learning was measured as conditioned approach (number of port entries made into each port while any of the CSs were on) and approach accuracy (fraction of entries that were into the paired port while a given CS was on). After the final day of training, mice underwent one 14 hour conditioned reinforcement test during which each entry into a port triggered a 2.5 s long presentation of the CS originally paired with that port, but no US was delivered. After the conditioned reinforcement test, mice also underwent one 14 hour primary reinforcement test during which each entry into a port triggered a 2.5 s long presentation of the US originally paired with that port, but no CS was presented. One mouse was run through the primary reinforcement test a second time because its optical patch cord came off partway through the first attempt.

### Statistical analysis

Statistical analyses were performed using R and GraphPad prism. Experiments in a two-factor design were analyzed using two-way ANOVAs with Geisser-Greenhouse corrections where possible or using mixed-effects models with subject as a random factor, followed by Holm-Sidak post-hoc tests. Experiments in a one-factor design were analyzed using parametric tests (t-tests or one-way ANOVAs with Holm-Sidak post hoc tests) unless the data failed to meet the equal variance assumption (Barlett’s test) in which case nonparametric methods were used (Kruskal-Wallis or Friedman’s tests followed by Dunn’s post-hoc tests). Paired or repeated measures comparisons were used where appropriate. Where tests were made against a hypothetical mean, one-sample t-tests were used followed by Bonferroni’s method to correct for multiple comparisons if multiple groups were tested. All hypotheses tested were two-tailed, and data are shown as mean +/- s.e.m.

## ACKNOWLEDGMENTS

We thank Stephan Lammel, Liqun Luo, Lisa Giocomo, Gregory Scherrer, and members of the Malenka lab for discussions, Jayashri Viswanathan for help with histology and microscopy, and the Stanford Gene Vector and Virus Core for reagents. This work was supported by philanthropic funds donated to the Nancy Pritzker Laboratory at Stanford University. M.B.P. was supported by NIH grant K99DA056573. N.E. was supported by NIH grant K08MH123791, a Brain & Behavior Research Foundation Young Investigator Grant, a Burroughs Wellcome Fund Career Award for Medical Scientists, a Stanford NeuroChoice Initiative Pilot Award, and a Simons Foundation Bridge to Independence Award. D.F.C.P. was supported by an NSF Graduate Research Fellowship and an HHMI Gilliam Fellowship for Advanced Study (with R.C.M.).

## AUTHOR CONTRIBUTIONS

D.F.C.P. and R.C.M. conceived the study and designed the experiments with input from M.B.P. and N.E. Mouse line characterization experiments were performed by D.F.C.P and M.G. Anatomical tracing experiments were performed by D.F.C.P., and M.G. B.S.B. and N.E. built the fiber photometry rig and wrote analysis software. Fiber photometry experiments were performed by D.F.C.P. Optogenetic experiments were performed by D.F.C.P and M.B.P. D.F.C.P. analyzed the data and interpreted it together with M.B.P., N.E., and R.C.M. The manuscript was written by D.F.C.P. and R.C.M. and edited by all authors.

## DATA AVAILABILITY

The datasets generated and analyzed during this study are available from the corresponding author upon reasonable request

## CODE AVAILABILITY

Code used for data processing and analysis is available from the corresponding author upon reasonable request.

## CORRESPONDENCE

Correspondence and requests for materials should be addressed to Robert C. Malenka.

## EXTENDED DATA FIGURES AND FIGURE LEGENDS

**Extended Data Fig. 1:**
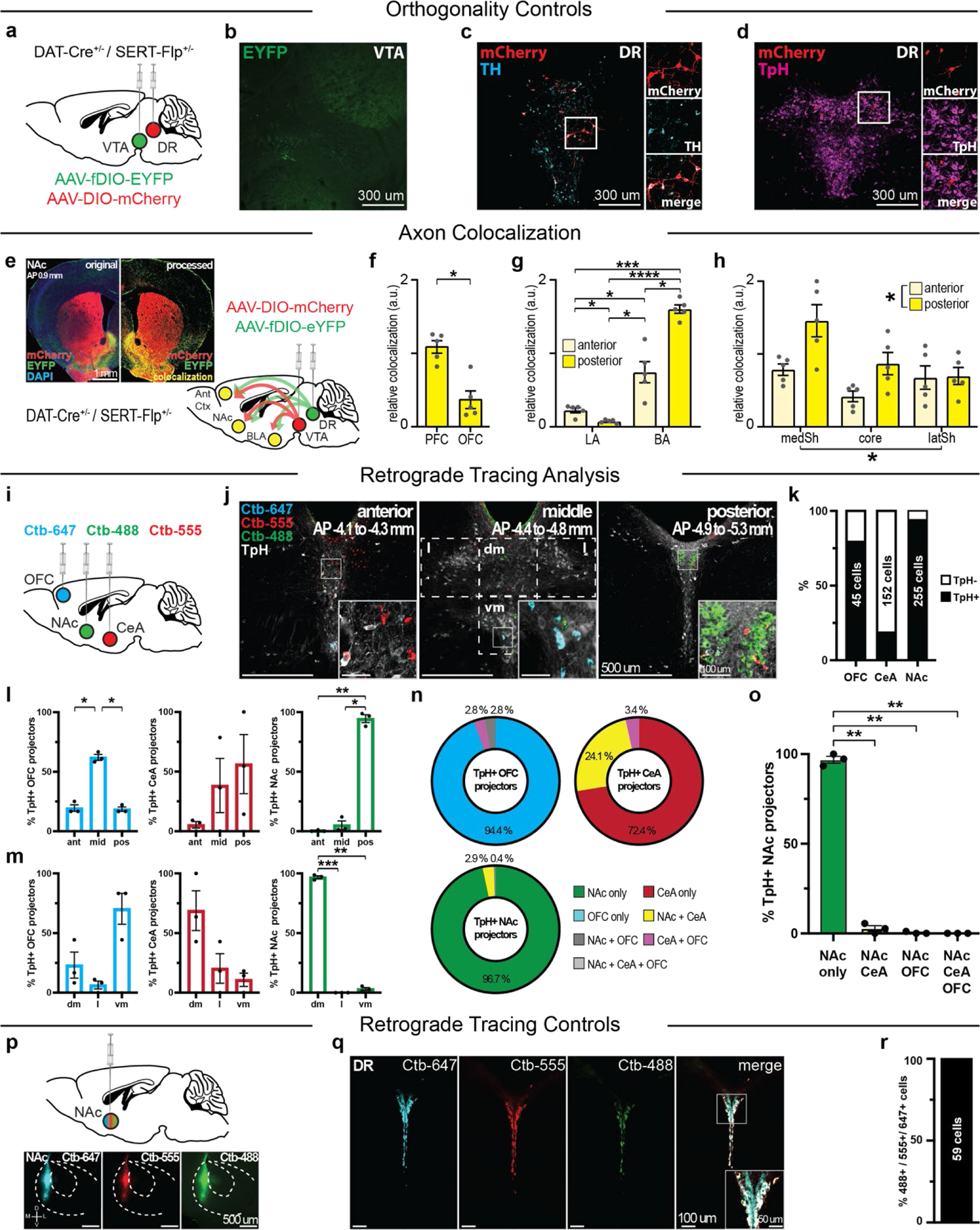
DAT-Cre^+/-^ / SERT-Flp^+/-^ mouse line characterization controls and additional anatomical tracing data. **a**, Viral strategy to validate orthogonality of genetic access to VTA^DA^ and DR^5HT^ neurons in DAT-Cre^+/-^ / SERT-Flp^+/-^ mice. **b**, Example image showing negligible Flp-dependent EYFP expression in the VTA. **c-d**, Example images of the DR showing Cre-dependent mCherry expression is restricted to DR^DA^ neurons (**c**) and is not observed in DR^5H^^T^ neurons (**d**). **e**, Viral strategy for VTA^DA^ and DR^5HT^ axon tracing experiments (right), and example raw and processed axon tracing images (left). **f-h**, Colocalization between VTA^DA^ and DR^5HT^ axons was heterogeneous across subregions of limbic structures (**f**, Ant Ctx, paired t-test: t_(4)_=4.233, P=0.0134; **g**, BLA, two-way RM ANOVA: subregion x AP interaction, F_(1.000,4.000)_=43.41, P=0.0027; Holm-Sidak tests: anterior LA vs posterior LA, P=0.0318; anterior LA vs anterior BA, P=0.0318; anterior LA vs posterior BA, P=0.0002; posterior LA vs anterior BA, P=0.0318; posterior LA vs posterior BA, P<0.0001; anterior BA vs posterior BA, P=0.0315; **h**, NAc, two-way RM ANOVA: main effect of subregion, F_(1.562,6.249)_=6.132, P=0.0380; main effect of AP, F_(1.000,4.000)_=8.086, P=0.0467; subregion x AP interaction, F_(1.130,4.518)_=2.589, P=0.1761; **f-h**, n=5 mice). **i**, Surgical strategy for retrograde labeling of projection-defined DR^5HT^ subsystems. **j**, Example images of the DR showing retrogradely labeled neurons (posterior DR image reproduced here from Fig. 1m for comparison). **k**, Percentage of OFC-, CeA-, and NAc- projecting DR neurons that are TpH+ (n=3 mice). **l-m**, Distributions of OFC- (left), CeA- (center), and NAc- (right) projecting DR^5HT^ neurons across the DR’s anteroposterior axis (**l**, OFC, one-way RM ANOVA: F_(1.676,3.352)_=72.17, P=0.0019; Holm-Sidak tests: ant vs mid, P=0.0259; ant vs pos, P=0.8376; mid vs pos, P=0.0175; CeA one-way RM ANOVA: F_(1.009,2.018)_=1.127, P=0.3923, n=3 mice; NAc, one-way RM ANOVA: F_(1.015,2.030)_=247.9, P=0.0038; Holm-Sidak tests: ant vs mid, P=0.3141; ant vs pos, P=0.0034; mid vs pos, P=0.0111; n=3 mice) and the subregions defined in **j** (**m**, OFC, one-way RM ANOVA: F_(1.070,2.141)_=7.361, P=0.1059; CeA one-way RM ANOVA: F_(1.075,2.151)_=4.127, P=0.1705; NAc, one-way RM ANOVA: F_(1.000,2.000)_=1985, P=0.0005; Holm-Sidak tests: dm vs l, P=0.0005; dm vs vm, P=0.0014; l vs vm, P=0.1243; n=3 mice). **n**, Pie graphs showing the fraction of OFC- (top left), CeA- (top right), and NAc- (bottom left) projecting DR^5HT^ neurons that send axon collaterals to the other two projection targets. **o**, NAc- projecting DR^5HT^ neurons constitute a third, projection-defined DR^5HT^ subpopulation distinct from the CeA- and OFC- projecting subsystems (one-way RM ANOVA: F_(1.012,2.025)_=1090, P=0.0009, n=3 mice; Holm- Sidak tests: NAc only vs NAc+CeA, P=0.0055; NAc only vs NAc+OFC, P=0.0027; NAc only vs NAc+CeA+OFC, P=0.0023; all other comparisons p>0.05; n=3 mice). **p**, Injection strategy (top) and example injection site images (bottom) for retrograde tracing control experiments. **q-r**, when all three retrograde tracers were injected together into the same target structure, the vast majority of labeled cells in the DR were positive for all three tracers (n=59 cells from 2 mice) confirming that lack of colocalization observed in other experiments was not due to technical limitations of this approach.

**Extended Data Fig. 2:**
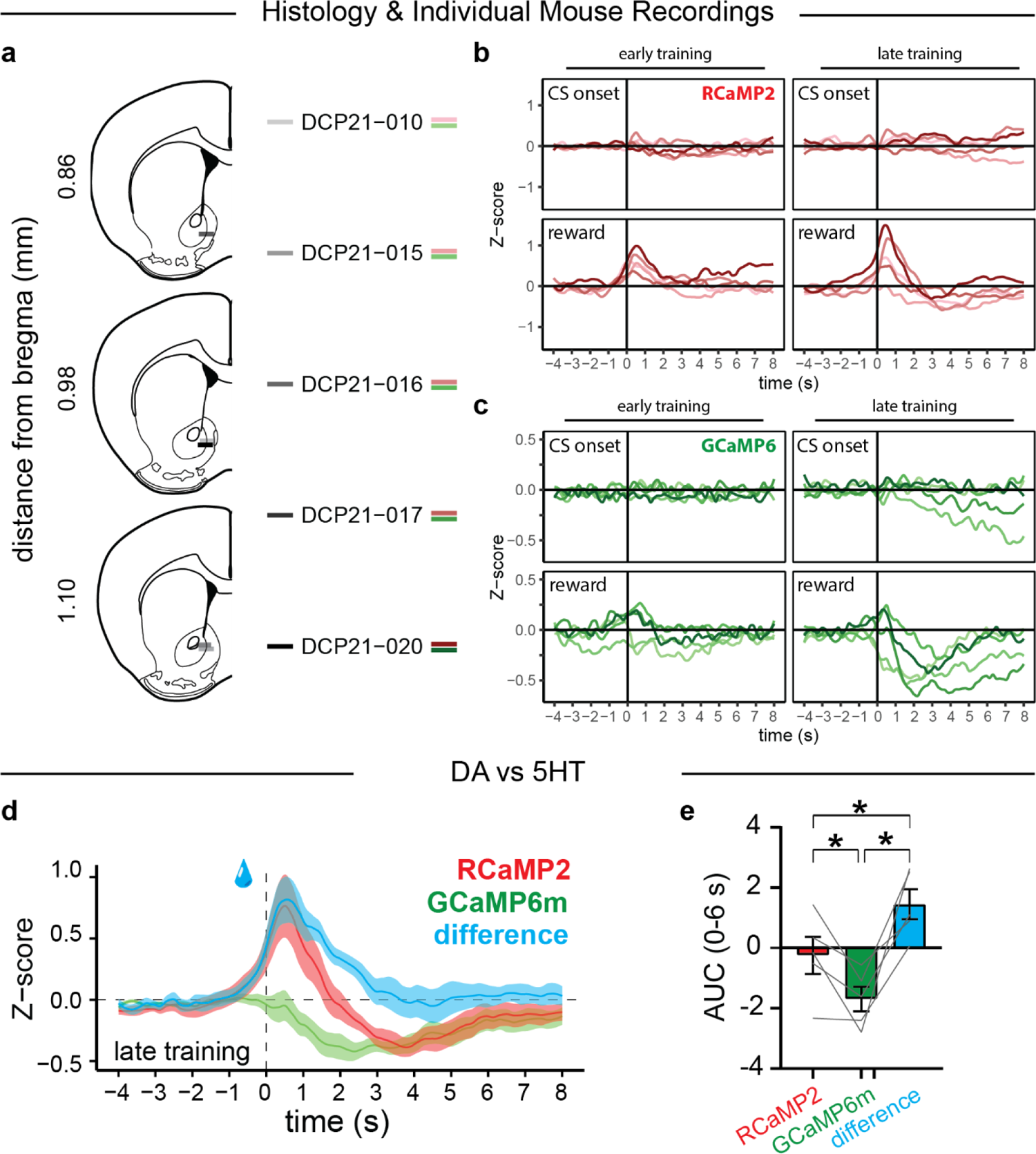
Histology and additional analyses for DA and 5HT photometry experiments **a**, Optical fiber tip placements for the mice used in the experiments in axon photometry experiments. **b-c**, RCaMP2 (**b**, VTA^DA^ axons) and GCaMP6 (**c**, DR^5HT^ axons) recordings for individual mice in response to CS-onset (top) and reward consumption (bottom) early (left) and late (right) in training. **d**, Average GCaMP6, RCaMP2, and RCaMP2-GCaMP6 (DIFF) photometry traces aligned to reward consumption. **e**, Area under the curve quantification for the traces in **d** (one-way RM ANOVA: F_(1.498,5.991)_=17.29, P=0.0042, n=5 mice; Holm-Sidak tests: RCaMP vs GCaMP, P=0.0428; RCaMP vs DIFF, P=0.0287; GCaMP vs DIFF, P=0.0275).

**Extended Data Fig. 3:**
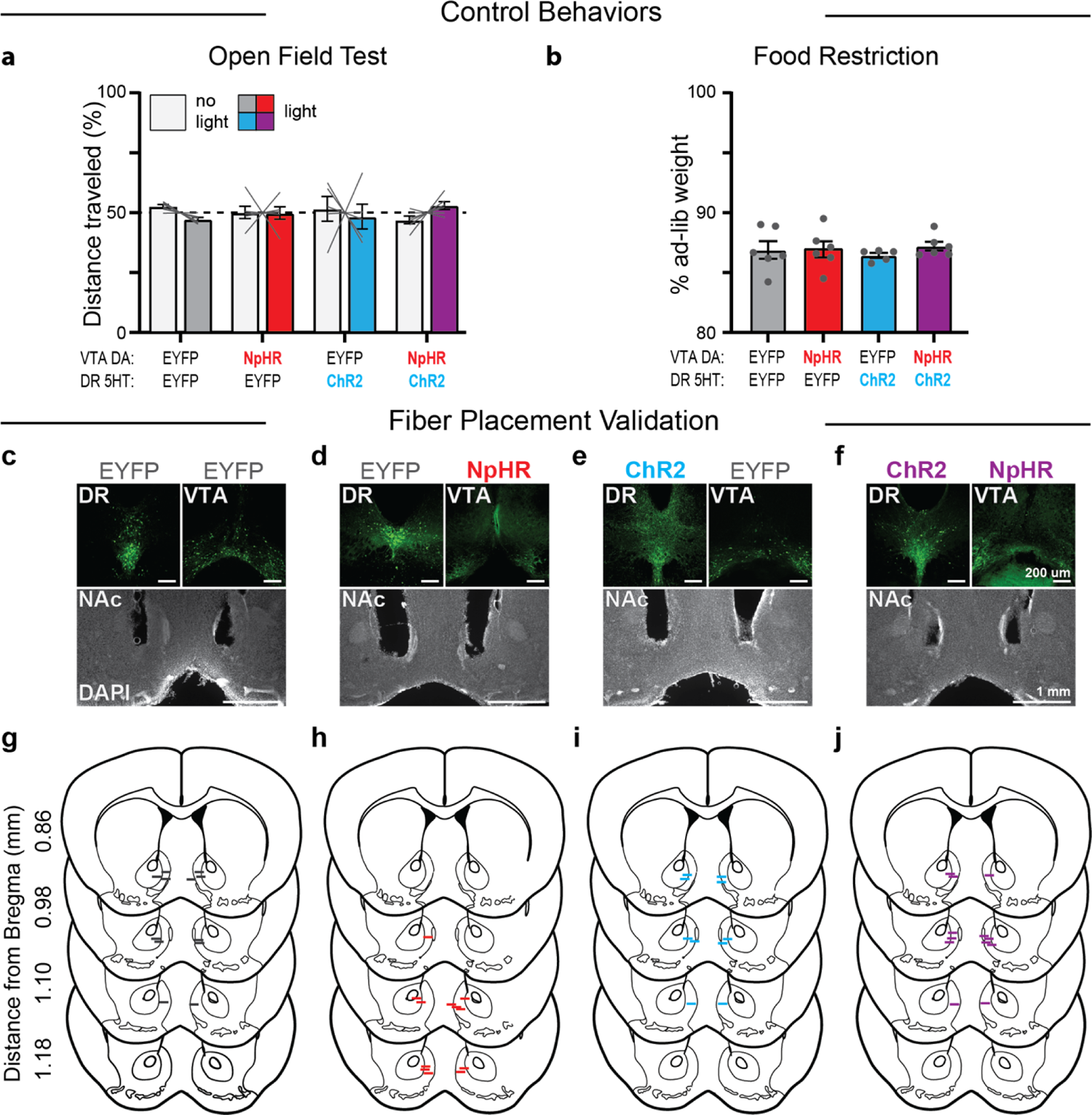
Control assays and histology for loss-of-function experiments **a**, Loss-of-function manipulations did not affect locomotion in the open field test (mixed-effects model: fixed effect of virus, F_(3,19)_<0.0001, P>0.9999; fixed effect of light, F_(1,19)_=0.1364, P=0.7159; virus x light interaction F_(3,19)_=1.714, P=0.1981, n=5-6 mice/group). **b**, All groups were equally food restricted during the appetitive conditioning task (one-way ANOVA: F_(3,19)_=0.2980, P=0.8264, n=5-6 mice/group). **c-f**, Example images of the injection sites (DR, top left; VTA, top right) and optical fiber implantation sites (bottom) for the EYFP/EYFP (**c**), EYFP/NpHR (**d**), ChR2/EYFP (**e**), and ChR2/NpHR (**f**) groups. **g-j**, Optical fiber tip placements for mice in the EYFP/EYFP (**g**), EYFP/NpHR (**h**), ChR2/EYFP (**i**), and ChR2/NpHR (**j**) groups.

**Extended Data Fig. 4:**
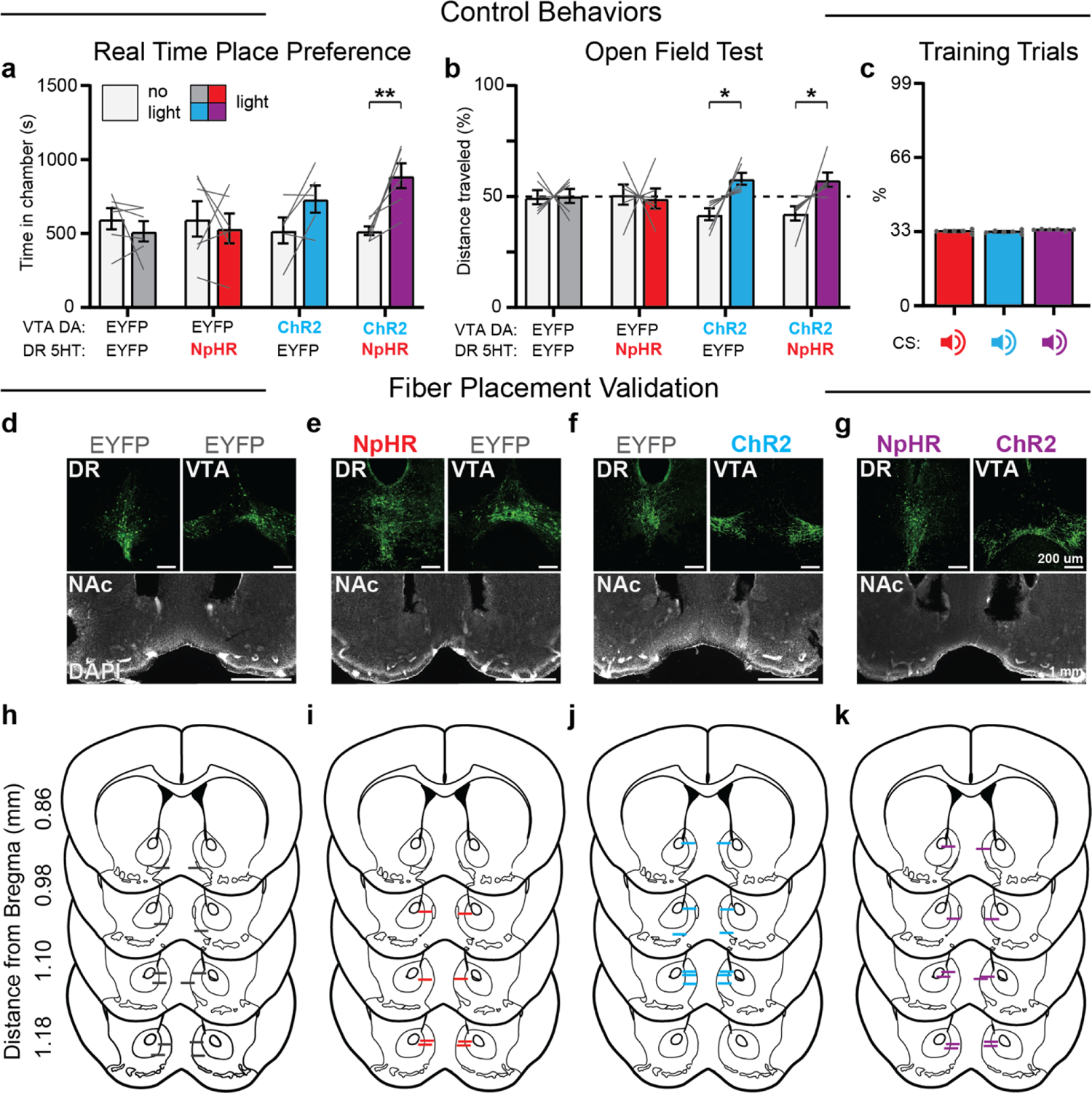
Control assays and histology for gain-of-function experiments **a**, VTA^DA^ excitation delivered together with DR^5HT^ inhibition together was rewarding in the real-time place preference test (two-way RM ANOVA: virus x light interaction, F_(3,19)_=4.539, P=0.0146, n=5-6 mice/group; Holm-Sidak tests, no light vs light: EYFP/EYFP, P=0.6685; EYFP/NpHR, P=0.6685; ChR2/EYFP, P=0.2114; ChR2/NpHR, P=0.0077). One mouse in the ChR2/EYFP group was excluded because it only explored one side of the behavioral apparatus during the test. **b**, VTA^DA^ excitation alone and VTA^DA^ excitation delivered together with DR^5HT^ inhibition increased locomotion in the open field test (mixed-effects model: virus x light interaction F_(3,20)_=3.721, P=0.0283, n=6 mice/group; Holm-Sidak tests, no light vs light: EYFP/EYFP, P=0.9219; EYFP/NpHR, P=0.9219; ChR2/EYFP, P=0.0145; ChR2/NpHR, P=0.0165). **c**, In the optogenetic conditioning experiment shown in Fig. 4g-j, mice were exposed to equal numbers of each trial type (one-way RM ANOVA: F_(1.302,6.512)_=1.1910, P=0.2182, n=6 mice). **d-g**, Example images of the injection sites (DR, top left; VTA, top right) and optical fiber implantation sites (bottom) for the EYFP/EYFP (**d**), NpHR/EYFP (**e**), EYFP/ChR2 (**f**), and NpHR/ChR2 (**g**) groups. **h-k**, Optical fiber tip placements for mice in the EYFP/EYFP (**h**), NpHR/EYFP (**i**), EYFP/ChR2 (**j**), and NpHR/ChR2 (**k**) groups. Two mice in the NpHR/EYFP group died before their brains could be collected for histology.

